## Supplementary material for "HippoUnit: A software tool for the automated testing and systematic comparison of detailed models of hippocampal neurons based on electrophysiological data": S1 Appendix

### Example of running the SomaticFeaturesTest of HippoUnit using a Jupyter notebook

As the first step HippoUnit’s test classes and ModelLoader class, along with a few additional Python packages must be imported:

1. **from** **__future__** **import** print_function #needed only in Python 2
2. % matplotlib inline
4. **from** hippounit.utils **import** ModelLoader
5. **from** hippounit **import** tests
7. **import** pkg_resources
8. **import** json
9. **import** collections
10. **import** numpy

Then the path to external mechanisms used by the Neuron implementation of the model (NMODL files) needs to be provided, which will be an argument to the ModelLoader class, so that the NMODL files can be compiled when the ModelLoader class is instantiated (if they are not compiled yet). Next, the variables related to the model are set. The initial voltage (v_init) and temperature (celsius) values specific to the model need to be set; otherwise, the default value in the corresponding capability method of the ModelLoader will be used. Setting the cvode_active boolean parameter to True or False, the user can decide to run the simulations using variable or fixed time step, respectively.

11. # path to NMODL files
12. mod_files_path = "/home/saray/published_models/Ca1_Bianchi_2012/experiment/"
14. #all the outputs will be saved here. It will be an argument to the test.
15. base_directory = '/mnt/csoport31-2/Modellezo_csapat/Sara/published_models_validation_results/'
17. #Load cell model
18. model = ModelLoader(mod_files_path = mod_files_path )
20. # outputs will be saved in subfolders named like this:
21. model.name="Bianchi_et_al_2012"
23. # path to hoc file
24. # the model must not display any GUI!!
25. model.hocpath = "/home/saray/published_models/Ca1_Bianchi_2012/experiment/main_model.hoc"
27. # If the hoc file doesn't contain a template, this must be None (the default value is None)
28. model.template_name = None
30. # model.SomaSecList_name should be None, if there is no Section List in the model for the soma, or if the name of the soma section is given by setting model.soma (the default value is None)
31. model.SomaSecList_name = None
32. # if the soma is not in a section list or to use a specific somatic section, add its name here:
33. model.soma = 'soma[0]'
35. # For the PSP Attenuation Test, and Back-propagating AP Test a section list containing the trunk sections is needed
36. model.TrunkSecList_name = 'apical_trunk_list'
37. # For the Oblique Integration Test a section list containing the oblique dendritic sections is needed
38. model.ObliqueSecList_name = 'oblique_dendrites'
39. # This will be argument to those tests, where dendritic locatins are selected according to distances. If not set, the end of the above given soma section will be used as reference point for distance determination
40. trunk_origin = ['soma[0]', 1]
42. model.v_init = -70
43. model.celsius = 34
45. # It is possible to run the simulations using variable time step (default for this is False)
46. model.cvode_active = True

The target experimental data and the configuration file are loaded from the JSON files to the *observation* and *config* dictionaries, which are arguments of the test class:

47. # Load target data
48. with open('/home/saray/target_features/feat_CA1_pyr_cACpyr_more_features.json') as f:
49. observation = json.load(f, object_pairs_hook=collections.OrderedDict)
51. # Load stimuli file
52. ttype = "CA1_pyr_cACpyr"
54. stim_file = pkg_resources.resource_filename("hippounit", "tests/stimuli/somafeat_stim/stim_" + ttype + ".json")
55. with open(stim_file, 'r') as f:
56. config = json.load(f, object_pairs_hook=collections.OrderedDict)

Then the test class is instantiated, and its judge() function (inherited from SciUnit) is called to run the test. The number of parallel processes to be used can be controlled by the user by setting the test.npool parameter.

57. # Instantiate test class
58. test = tests.SomaticFeaturesTest(observation=observation, config=config, force_run=False, show_plot=True, save_all = True, base_directory=base_directory)
60. # test.specify_data_set is added to the name of the subdirectory (somaticfeat), so test runs using different data sets can be saved into different directories
61. test.specify_data_set = 'UCL_data'
63. # Number of parallel processes
64. test.npool = 30
65. #Run the test
66. score = test.judge(model)
67. #Summarize and print the score achieved by the model on the test using SciUnit's summarize function
68. score.summarize()

For further details on how to run the different tests of HippoUnit for the different models, see the Jupyter Notebooks available here: <https://github.com/KaliLab/HippoUnit_demo/tree/master/jupyter_notebooks>.
